# Coherent thalamic inputs organize head direction signal in the medial entorhinal cortex

**DOI:** 10.1101/2025.05.14.654107

**Authors:** Gilberto R. Vite, Michelle Ding, Adrien Peyrache

## Abstract

Successful navigation relies on signals that remain stable despite environmental changes. This stability can arise by constraining neuronal activity to low-dimensional subspaces. During sleep, when external input is reduced, pairwise coordination persists in the spatial navigation system, as observed for head-direction (HD) cells of the anterodorsal nucleus (ADn) and grid cells of the medial entorhinal cortex (MEC). As ADn is crucial for spatial representation in the MEC, we hypothesized that coherent HD input underlies MEC organization. To test this, we performed simultaneous recordings in the ADn and MEC during wakefulness and sleep. We found that HD cell pairs maintained stable coordination across both states, and that MEC cell coordination was partly driven by common inputs from ADn HD cells. These results suggest that MEC activity is shaped, in part, by coherent thalamic HD signals, supporting stable network organization.

## Introduction

Animals use a ‘cognitive map’ to flexibly navigate their environment, which is supported by diverse functional cell types in the hippocampal formation and associated structures (McNaughton et al., 2006; O’Keefe & Nadel, 1978; Tolman, 1948). Head-direction cells form an important backbone of these representations (McNaughton et al., 2006; Taube et al., 1990) and are critical for the emergence of higher-level spatial signals in the medial entorhinal cortex (MEC), especially grid cells (Burak & Fiete, 2009; Couey et al., 2013; Fuhs & Touretzky, 2006; Hafting et al., 2005; Winter et al., 2015). The anterodorsal nucleus (ADn) of the thalamus is the main source of the HD signal to the cortex, and a high number of HD cells are found across the subicular complex and in the MEC (Boccara et al., 2010; Giocomo et al., 2014; Obenhaus et al., 2022).

Importantly, HD cells are believed to form a continuous attractor network, whereby cellular and circuit properties constrain the states of the neuronal population to a functional ring (Knierim & Zhang, 2012; McNaughton et al., 2006; Redish et al., 1996; Sharp et al., 2001; W. E. Skaggs et al., 1995). In support of this view, pairwise correlation is preserved among HD cells during sleep, when sensory inputs are virtually absent, not only in the ADn (Chaudhuri et al., 2019; Peyrache et al., 2015) but also in the associated cortical areas (Duszkiewicz et al., 2024; Gardner et al., 2019; Peyrache et al., 2015). However, it is unclear whether these co-activity patterns arise locally or are orchestrated by common inputs from the ADn.

The ADn projects mainly to the postsubiculum (PoSub) as well as to the retrosplenial cortex (RSC) (Peyrache et al., 2015; Shibata, 1993; Swanson & Cowan, 1977; van der Goes et al., 2024; Van Groen & Wyss, 1995). In turn, both the PoSub and the RSC project to the MEC (Kononenko & Witter, 2012; Preston-Ferrer et al., 2016; Simonsen et al., 2022; Tukker et al., 2015). During sleep, HD cells in the ADn-PoSub network maintain their coordination (Peyrache et al., 2015), and the PoSub thus provides the MEC with a coherent HD signal across all brain states. The ADn can influence MEC activity indirectly, through polysynaptic pathways. Therefore, the observation that in the MEC both HD cells and grid cells (Gardner et al., 2019; Trettel et al., 2019) maintain their mutual coupling during sleep suggests a local coordination of population activity. As such, not only HD cells but also grid cells would form local, independent attractors in the MEC that would be weakly coupled to other spatially modulated cell populations in associated brain areas. Another possibility is that the highly coherent HD signal conveyed by the ADn exerts strong control over the entire cortical navigation system, as suggested by the widespread influence of ADn activity on the excitability of posterior cortical areas (Gent et al., 2018).

In the present work, to address the question of whether pairwise coupling in the MEC arises from local interaction or is controlled by indirect inputs from the ADn, we performed recordings of neuronal ensembles in the ADn and the MEC using high-density silicon probes during freely moving exploration and natural sleep. We found that the HD signal remains in register across brain states and that pairwise coupling of MEC HD cells result in large part from common inputs, indirectly coordinated by the ADn.

## Results

### HD cells in ADn and MEC are coupled during sleep

To test whether neuronal ensembles remain coordinated in the ADn-MEC network across wake and sleep states, we implanted six freely moving mice with silicon probes and recorded 213 anterior thalamic neurons and 1004 MEC neurons (Figure 1A-D). Among these neurons, 88 thalamic cells were classified as HD cells, likely from the ADn, and 284 in the MEC were classified as HD cells based on HD information and stability. As expected, during exploratory behavior, the activity of ADn and MEC HD cells closely followed the actual head direction of the animal (Figure 1E). We used a Bayesian decoder (Methods) to extract the internal head direction of the animal from ADn and MEC spiking activity – only sessions where both structures had at least 4 HD neurons were considered for estimating the error (n = 9 sessions from 6 animals) with decoding error below chance level (obtained from the time-inverted MEC spikes) during the wake epoch (Figure 1E-F). During SWS and REM sleep, we observed that MEC was significantly aligned to ADn activity when compared to the control condition (Figure 1G), indicating that ADn and MEC keep coherent representation at the population level across sleep states, in the relative absence of incoming sensory information.

**Figure 1.**
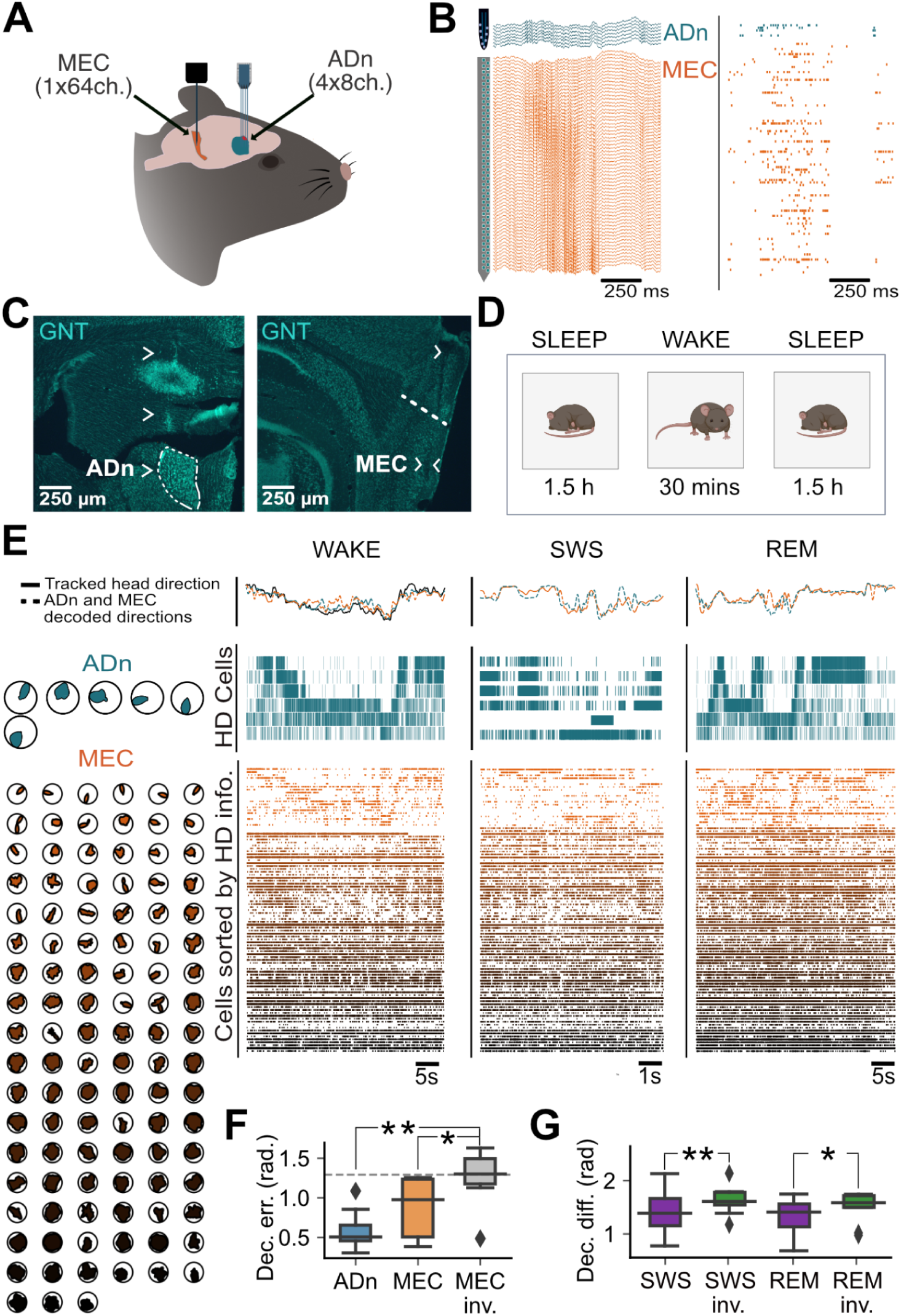
Coherent HD signal in the ADn and the MEC during wakefulness and sleep. (A) Schematic of the electrophysiological implant showing silicon probes targeting the anterodorsal thalamic nucleus (ADn; 32-channel probe) and the medial entorhinal cortex (MEC; 64-channel probe). (B) Traces of local field potentials (left) and spike raster plots (right) from ADn (blue) and MEC (orange) during SWS. (C) Histological sections showing probe placements in ADn and MEC using green NeuroTrace (GNT) Nissl stain. Arrowheads indicate electrode tracks. (D) Recording protocol consisting of a sleep–wake–sleep sequence. (E) Left: tuning curves of simultaneously recorded HD cells in ADn (top, blue, *n* = 6) and MEC (bottom, orange, *n* = 105). Right: raster plots of neuronal activity. MEC cells are sorted by HD information (HD cells are shown at the top, further sorted by preferred direction). Solid and dashed lines indicate the animal’s actual and decoded head directions, respectively. (F) Decoding error from ADn and MEC sessions relative to the animal’s head direction. The dotted line indicates chance level (based on time-inverted spike trains; see Methods). (G) Decoding differences between ADn and MEC (purple), and between ADn and time-inverted MEC HD cell spikes (green). **p < 0.01, *p < 0.05, Wilcoxon signed-rank test.

### Brain state independent coupling of ADn-MEC HD cells

HD cells in ADn and MEC maintain their pairwise correlation across behavioral states (Gardner et al., 2019; Peyrache et al., 2015). To determine whether HD cells were locally or globally coordinated, we evaluated pairwise coupling between the ADn and MEC. To achieve this, and considering the high degree of coupling among some cortical neurons (Okun et al., 2015), we used a Generalized Linear Model (GLM) to quantify the coupling between individual neuron pairs, while accounting for shared variability by regressing out the influence of local population activity (Garner et al;, 2019; Duszkiewicz et al., 2024) (see Methods).

While ADn HD cells are overall homogeneous (differing only in their peak firing rate), MEC HD cells are more diverse (Giocomo et al., 2014) and we further classified them based on the width of their tuning curve between broad (n = 204) and sharp (n = 80). Differences in HD tuning in MEC are not just a quantitative feature, since evidence shows that sharp and broad HD cells have different connectivity, suggesting they form separate subnetworks (Zutshi et al., 2018). For the sharp MEC HD cells, we observed strong coupling between ADn and MEC HD cells across wake and sleep (Figure 2A,B). Furthermore, the strength of this coupling depended on the pairwise angular offset in preferred directions. Specifically, pairs with small angular differences in their preferred firing directions during wakefulness exhibited positive coupling during sleep, while pairs with large angular differences were negatively coupled (Figure 2C). Importantly, the correlation of coupling with angular offset was qualitatively similar during wakefulness and sleep. Furthermore, We observed a time compression of the neuronal dynamics during SWS, consistent with previous reports on place cells, grid cells and HD cells (Gardner et al., 2019; Lee & Wilson, 2002; Nádasdy et al., 1999; Peyrache et al., 2015; Trettel et al., 2019). During sleep, the coupling of broad MEC HD cells with ADn was also correlated with their pairwise angular difference, but qualitatively less than sharp HD cells (Supplementary Figure 2). Overall, these observations suggest a global coordination of sharply tuned HD cells, while broadly tuned HD neurons may participate in more independent subnetworks. In the following analyses, we will focus on sharp MEC HD.

**Figure 2.**
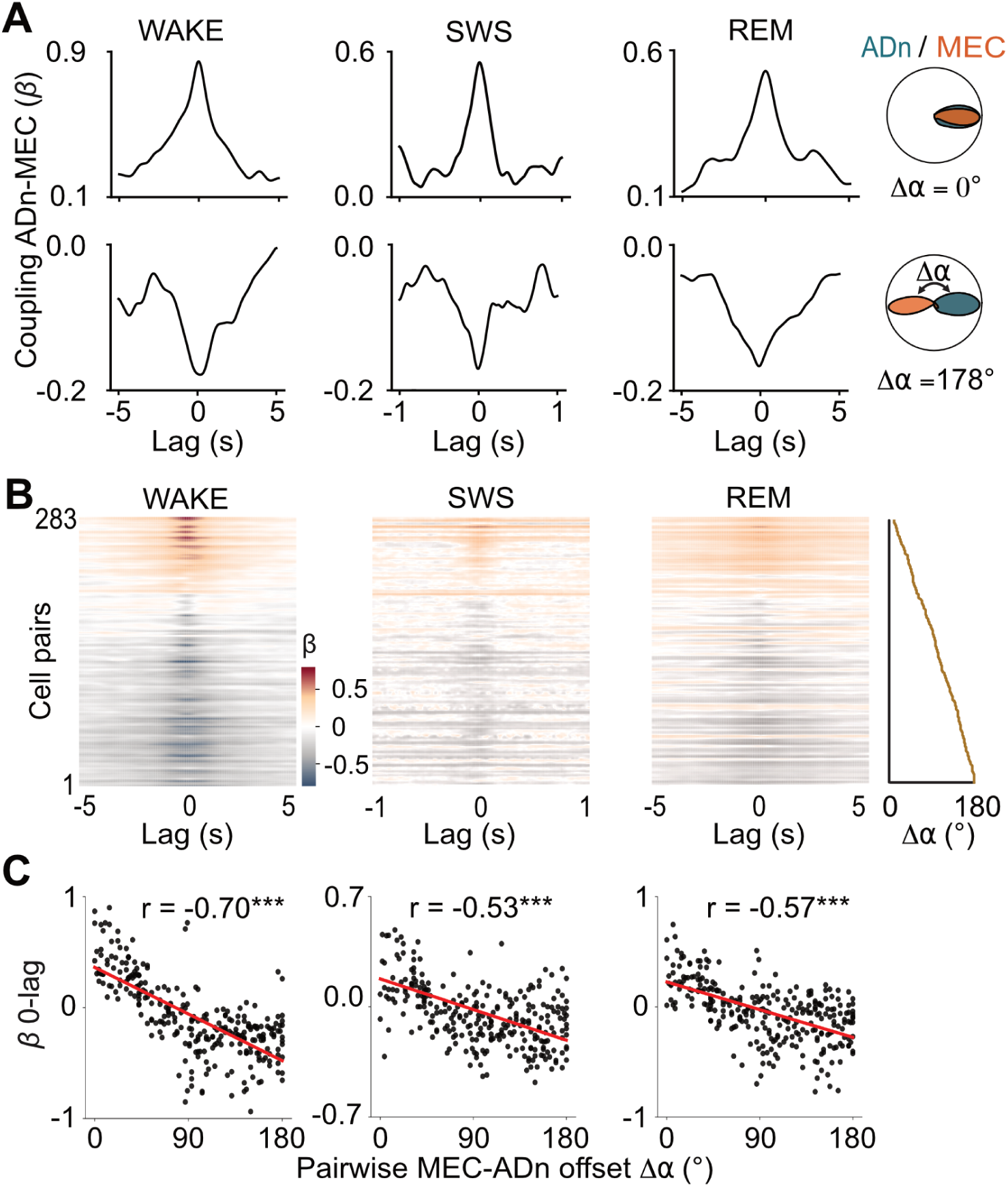
Pairwise correlation of ADn-MEC HD cell pairs is consistent across states. (A) HD tuning curves of two representative ADn–MEC cell pairs. Top: example pair with similar preferred directions (left), and the corresponding time-resolved pairwise coupling across wakefulness, SWS, and REM sleep. GLM beta coefficients were computed independently at each time lag. Bottom: same as above for a cell pair with opposite preferred directions. (B) Time-resolved coupling of all ADn–sharply tuned MEC cell pairs recorded during wakefulness and sleep, sorted by angular offset (*n* = 283 ADn–MEC pairs). Color indicates coupling strength, from warm (positive) to dark (negative) values. (C) Pairwise coupling at 0-lag as a function of angular offset. Spearman correlation coefficients (*r*) are indicated at the top; *p* < 0.001 for all brain states.

HD cells in the MEC show different levels of modulation by the theta rhythm. Previous reports suggest that only theta-modulated cells exhibit attractor dynamics (Kornienko et al., 2018). In line with these findings, we classified our dataset into theta-modulated and non-theta-modulated cells (Supplementary Figure 2A), with non-theta-modulated cells forming a minority of neurons (around 10% of sharp MEC HD cells). We observed that the ADn coupling with MEC HD cells was generally weaker for non-theta-modulated cells, with this reduction being especially pronounced during SWS, as indicated by the lower Pearson correlation values of coupling at 0-lag versus the tuning curve offset (Supplementary Figure 2B). Overall, these findings support the idea of weakly coupled cell populations within the MEC, and theta-modulated sharp HD cells form a subnetwork strongly coupled to the ADn.

### MEC HD cells are specifically coupled with ADn

To determine whether ADn neurons were coupled to other functional cell classes in the MEC, we quantified the coupling between ADn neurons and five categories of MEC neurons: (1) sharply tuned HD cells, (2) grid cells, (3) speed cells, (4) conjunctive HD × grid cells, and (5) fast-spiking interneurons. Because both positively and negatively signed loadings can represent robust interactions (Qian et al., 2023), we used the absolute value of the zero-lag GLM coefficient as our index of coupling strength. MEC sharp HD cells exhibited the highest coupling with ADn (Figure 3) and, overall, the level of ADn-MEC coupling was correlated with the amount of HD information (i.e. sharpness of HD tuning curve) of the MEC cell. This pattern was consistent across wakefulness, SWS, and REM sleep.

**Figure 3.**
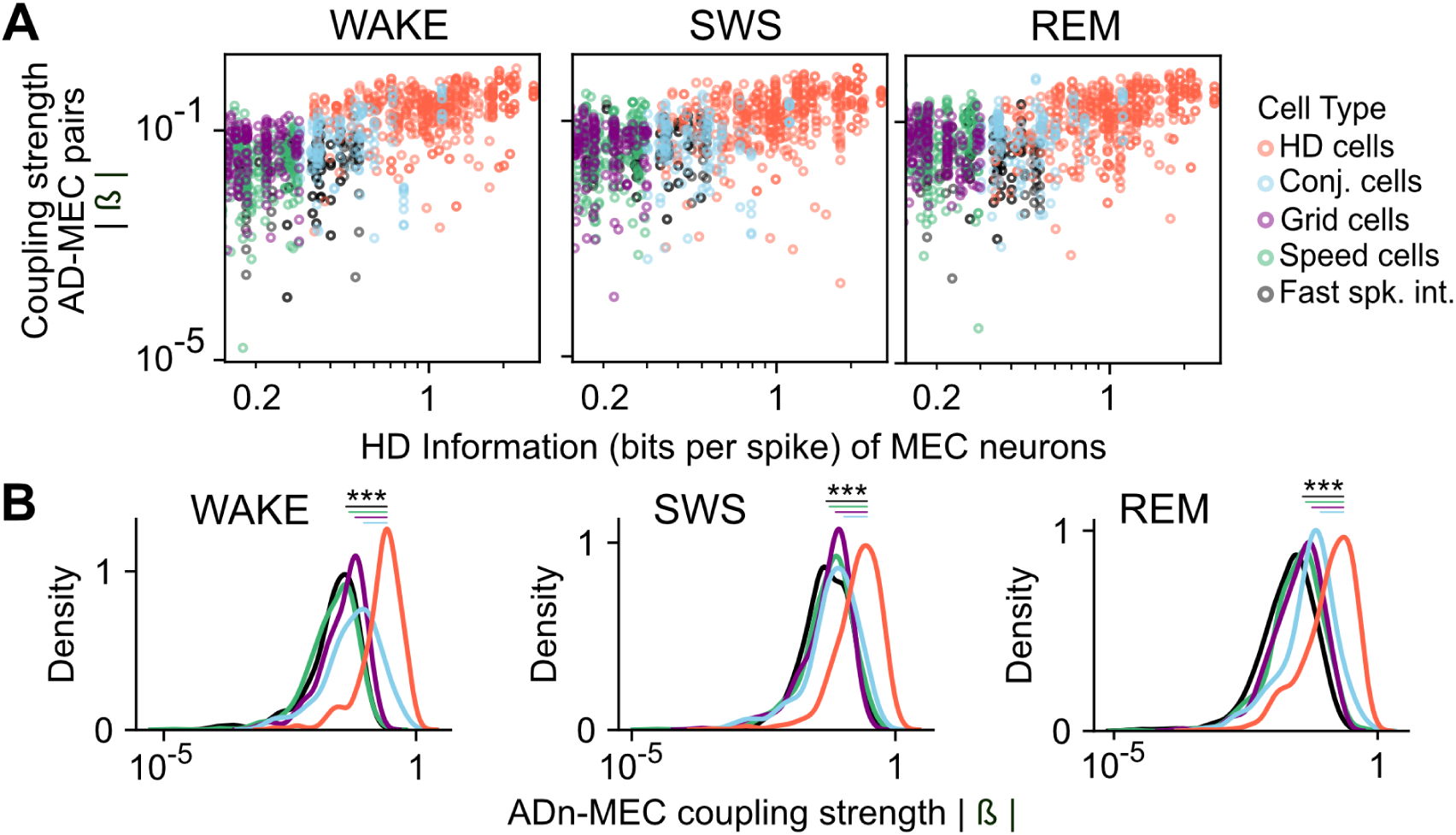
Sharply tuned MEC HD cells are distinctively coupled to the ADn HD signal. (A) ADn–MEC coupling strength (absolute GLM beta values at zero lag) plotted against the HD information of MEC neurons, color-coded by cell type. (B) Density distributions of coupling strength for all evaluated cell types. \*\*\**p* < 0.001, Bonferroni-corrected *p*-values from two-sample Kolmogorov–Smirnov tests.

### Pairwise coupling of sharp MEC HD cells depends on ADn input

MEC HD cells maintain their coupling during sleep (Gardner et al., 2019; Trettel et al., 2019) but does this coupling arise locally? To address this question, we first conditioned MEC-MEC coupling on ADn inputs. Specifically, we extended the GLM approach to triplets of cells. We regressed MEC HD cells activity using three predictors: the spiking activity of another MEC HD cell, the spiking activity of an ADn cell, and the overall MEC population activity. Finally, to assess the influence of ADn input on local MEC coordination, we determined the input “gain” of MEC-MEC coupling as the difference between the actual coupling and the coupling in a null model where ADn activity was shuffled.

We observed a significant gain for triplets of MEC–MEC–ADn neurons in which the ADn HD cells had the same preferred direction as at least one of the two MEC HD cells, during wakefulness and REM. In contrast, when the ADn’s tuning curve was opposite that of the two MEC cells, shuffling had minimal impact on MEC-MEC coupling (Figure 4A,B). To control for the directionality of the coupling, we conditioned the coupling of ADn–HD cell pairs on the firing of MEC HD cells. The coupling of AD HD cell pairs was unaffected by the firing of MEC cells (Figure 4C).

**Figure 4.**
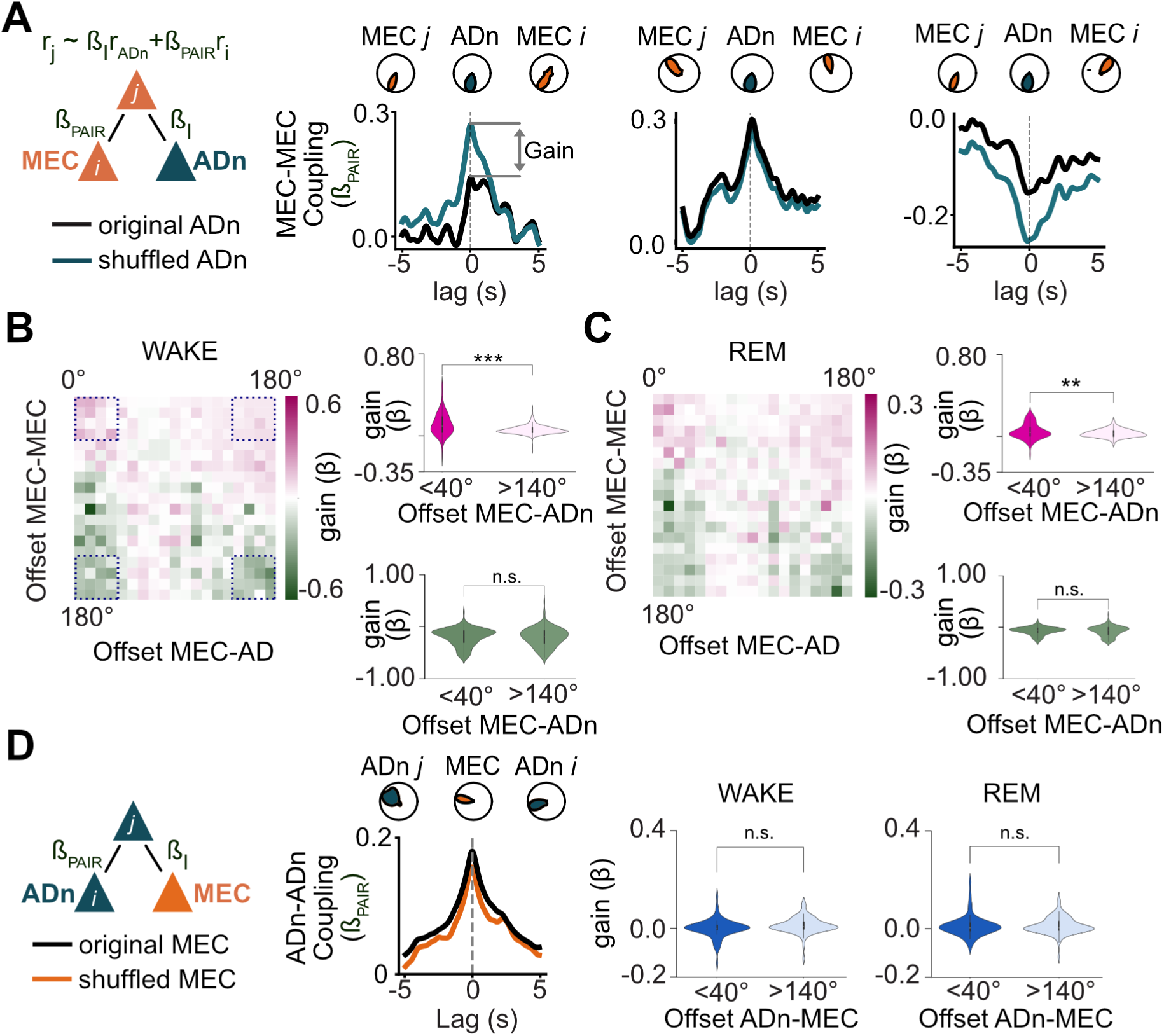
MEC HD cells pairwise coordination is explained by a common input. (A) Three examples of sharply tuned MEC HD cell activity modeled as a function of another MEC HD cell, an ADn HD cell, and the MEC population activity. *Gain* was defined as the difference between the original 0-lag MEC–MEC pairwise coupling coefficient and the corresponding coefficient obtained after shuffling the ADn spike train. (B) Left: coupling gain for all MEC–MEC–ADn HD cell triplets (*n* = 3320). Top right: coupling gains for triplets in which MEC cells had similar (left, *n* = 121) or opposite (right, *n* = 243) preferred directions relative to the ADn HD cell. Bottom right: same as top, but for triplets in which the ADn cell had a similar preferred direction to the target cell (left, *n* = 179) or the predictor MEC HD cell (right, *n* = 205). Note that in the latter case, the GLM returns similar results when the roles of the target and predictor MEC cells are swapped. (C) Same as (B), but during REM sleep. (D) Same as (B), but for ADn–ADn coupling in relation to MEC HD cell firing. \*\**p* < 0.001, \**p* < 0.01, Wilcoxon signed-rank test.

Overall, these observations suggest a primarily unilateral influence from ADn-HD cells onto MEC activity rather than mutual interdependence. These observations suggest that common ADn inputs at least partially account for co-firing of sharp MEC HD cells, even through a polysynaptic pathway, during wakefulness and REM sleep.

## Discussion

By recording neuronal ensembles simultaneously in the ADn and MEC, we found that both structures code for a common HD signal during sleep, when sensory inputs are largely absent. This suggests that the ADn–MEC circuit remains functionally coupled independently of ongoing sensory drive. Notably, this coupling was particularly strong for MEC HD cells that were sharply tuned and modulated by theta oscillations, whereas more broadly tuned and/or non-theta-modulated HD cells exhibited weaker coupling to the ADn. These findings point to the existence of distinct, weakly coupled subnetworks within the MEC, one of which, characterized by a sharp HD signal, appears to be organized by thalamic input, possibly via polysynaptic pathways.

A prevailing view of the MEC is that it exhibits attractor dynamics, in which population activity is constrained to low-dimensional substates. This view is supported by the coherent activity of grid and HD cells during sleep (Gardner et al., 2019, 2022; Trettel et al., 2019). Our findings show that the coupling of sharply tuned MEC HD cells is, in part, explained by inputs from the thalamus. This raises the possibility that these cells are less governed by local circuit dynamics and instead primarily relay the HD signal from the ADn.

The MEC contains a wide range of functionally defined cell types, including head-direction (HD), border, speed cells, and grid cells (Giocomo et al., 2014; Hafting et al., 2005; Kropff et al., 2015; Sargolini et al., 2006; Solstad et al., 2008). Rather than forming discrete categories, these cell types appear to exist along a continuum in feature space (Kropff et al., 2015; Obenhaus et al., 2022). Whether these functional correlates reflect distinct, weakly coupled subnetworks or distributed representations within a unified computational framework remains unclear. Supporting the former view, MEC HD cells have been shown to respond differently to changes in context; for example, non–theta-modulated HD cells often remap non-coherently across environments (Kornienko et al., 2018). Anatomical organization may also contribute to this functional heterogeneity. Grid cells are known to form modules along the dorsoventral axis (Stensola et al., 2012) and cluster anatomically (Gu et al., 2018). Similarly, HD tuning resolution varies along the dorsoventral axis in layers II-III in mice (layer III in rats) but remains consistent in deep layers (Giocomo et al., 2014). Our finding that sharply tuned MEC HD cells are preferentially coupled to ADn HD cells suggests that such subnetworks can also be defined by their coupling to specific external inputs.

Broadly tuned HD cells in the MEC are less correlated with ADn HD cells than their sharply tuned counterparts. Recent evidence suggests the existence of an internal directional signal that does not strictly follow allocentric head direction but instead alternates left and right of the current heading on alternating theta cycles (Vollan et al., 2025), producing an apparent broadening of tuning in HD space. This signal originates in the parasubiculum and remains coherent during sleep, pointing to a parallel circuit for directional coding that is only weakly dependent on the canonical HD system. Another potential source of directional input is the retrosplenial cortex (RSC), where directionally tuned neurons are more strongly driven by visual landmarks than by the core thalamocortical HD system (Jacob et al., 2017). Whether non–theta-modulated HD cells in the MEC are influenced by RSC or other visually anchored directional signals remains an open question.

The origin of a coherent HD signal during sleep remains unknown (Peyrache et al., 2015; Senzai & Scanziani, 2022) and likely varies across brain states. During REM sleep, changes in the HD signal correlate with eye movements, suggesting that the brainstem structures responsible for generating the HD signal during wakefulness remain active in this phase (Senzai & Scanziani, 2022). Our findings indicate that this signal is propagated upstream to the MEC. In contrast, the picture during SWS is more complex. ADn HD cells remain strongly organized during SWS (Peyrache et al., 2015) and have been shown to lead their cortical targets (Peyrache et al., 2017; Viejo & Peyrache, 2020), but it remains unclear whether the brainstem continues to generate the HD signal in this state. One possibility is that the ADn actively drives posterior cortical areas during SWS (Gent et al., 2018), possibly independently of brainstem inputs.

Together, these findings support a model in which the MEC comprises multiple, partially independent subnetworks, each shaped by distinct sources of input, including both long-range projections and local circuit mechanisms.

## Materials and Methods

### Animals

All animal experiments were approved and conducted in accordance with the guidelines of the Animal Care Committee at McGill University. Six adult male mice (approximately 45g each) were used in this experiment. These mice were from the first-generation of a cross between C57BL/GJ and FCB/NJ strains from Jackson Laboratories. We selected these mice because they are larger, less anxious and exhibit better running behaviour than those of the C57BL/GJ background (Sloin et al., 2022).

### Tissue processing

Anesthetized mice were perfused with a saline solution (0.9%) followed by cold paraformaldehyde in phosphate-buffered saline (PBS) at 4°C. Brains were post-fixed in the same PBS solution for 48 hours and then placed in the freezer for 24 hours. ADn and TRN sections were coronally sectioned at 40 µm using a freezing microtome. MEC sections were sagittally sectioned at 80 µm. Neurotrace Green Nissl stain was used for counterstaining in all sections.

### Implant surgeries

Animals were implanted under isoflurane anesthesia. Multi-electrode arrays (silicon probes) were implanted in the left hemisphere. The silicon probes were mounted on a nanodrive (Cambridge Neurotech) for recording neural activity and local field potentials (LFPs) in the anterior thalamus and MEC. The ADn was targeted with a Buszaki32 (Neuronexus; 160 µm² per site, ∼0.5-MΩ impedance), composed of four shanks (150 µm shank separation) and each with 8 recording sites. It was implanted at AP: −0.45; ML: −0.85; DV: −1.8, and it. To target the MEC, we used a single shank probe with 64 electrodes (H5, Cambridge Neurotech; 165 µm² per site, ∼0.1-MΩ impedance) in five animals,and a three shanks probe with 32 electrodes (Fb, Cambridge Neurotech; 165 µm² per site, ∼0.1-MΩ impedance) in one animal. Probes were implanted in the left hemisphere at AP: +0.2 from the transverse sinus; ML: −3.35; DV: −1.0, with a 7° angle.

### Recording procedures

Animals were habituated for one month to freely forage for food in an open field arena of 1m by 1m, (black walls with a white card as a cue in one of them). The lights of the room were dim and the arena was surrounded by gray curtains. Then we implanted these animals as described above. One week after the implant, animals were recorded according to the following protocol.

Sessions were divided in 3 blocks: sleep (∼120 min), wake (∼30 min), sleep (∼120 min). During the sleep session animals were put in the same cage where they are in the animal facilities. The animal position was tracked thanks to the detection of four reflective markers attached to the head cage of the implant by a system equipped with 11 infrared cameras using a sampling rate of 120 (fps). For the recording of the neurophysiological signals, silicon probes were connected to an Intan RHD2000 System, sampling frequency of 20kHz.

### Data processing

We utilized Kilosort 2.5 for the identification of single-unit activity. The resulting clusters of neurons were later refined manually using Klusters. We developed custom scripts in Python for analyzing neural activity, employing libraries such as NumPy for array operations, pandas for dataframe operations, pynapple (custom software for analyzing neural data originated at the Peyrache Lab (Viejo et al., 2023)) and SciPy (used for a broad range of computational analysis). For data visualization, we used the matplotlib and seaborn libraries. For the LFP of Figure 1E, the signal was downsampled to 1.25kHz using custom MATLAB scripts (Peyrache, 2015).

### Principal cell / interneuron classification

Cells were categorized based on their waveform characteristics and firing rates (Duszkiewicz et al., 2024). Fast-spiking (FS) interneurons, identified by their brief trough-to-peak duration (≤ 0.35ms) and high mean firing rates (> 10Hz), were distinguished from putative excitatory cells. The latter showed longer trough-to-peak durations (> 0.35 ms) and lower mean firing rates (< 10Hz).

### HD cell classification

Cells were classified as HD cells based on a strict criterion, considering head direction information, stability, and field size. HD information (Peyrache et al., 2015; W. Skaggs et al., 1992) helps identify cells whose firing rates are closely aligned with specific head orientations. HD information was quantified by measuring the information content (W. Skaggs et al., 1992):

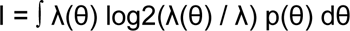

where:

– λ(θ) is the mean firing rate of the head direction cells at head direction θ,
– p(θ) is the probability density for the rat being at head direction θ,
– λ is the overall mean firing rate, computed as ∫ λ(θ) p(θ) dθ.

Cells with a HD information higher than 0.33 bits/spike were selected (Duszkiewicz et al., 2024). For cell stability assessment, we selected exploration epochs where the animal’s speed exceeded 3 cm/s. We divided the session into two halves, computed the tuning curves for each, and then calculated the correlation between these curves. Cells were considered stable if the correlation value exceeded 0.5 (p ≤ 0.05).

Finally, HD cells were classified as *sharp* if at least 50% of their firing was concentrated within ±45° of the circular mean of their tuning curve; otherwise, they were classified as *broad*.

### Speed cell classification

Following previous work (Kropff et al., 2015), we identified speed cells based on a speed score for each cell –defined as the Pearson product-moment correlation coefficient, ranging from −1 to 1, between the cell’s instantaneous firing rate and the rat’s instantaneous running speed. This analysis was restricted to exploration epochs where the speed of the animal was higher than 3 cm/s. Cells with a correlation value higher than 0.33 were considered speed cells.

### Grid cell classification

The classification of a cell as a grid cell was determined by its grid score (Sargolini et al., 2006). The grid score is derived from the spatial 2D autocorrelation of the cell’s firing rate map. The autocorrelogram is rotated every 6 degrees, and a circular area enclosing the inner peaks near the center—but excluding the central peak—is defined. This area is then correlated with its rotated version. The gridness score is calculated by subtracting the lowest correlation at 60 and 120 degrees from the highest correlation at 30, 90, and 150 degrees. Cells with a score above 0.2 are classified as grid cells. Some cells were discarded based on visual inspection.

### Sleep stages classification

Automated SleepScoreMaster algorithm and TheStateEditor software were employed for sleep scoring (Watson et al., 2016).

### Extracting the internal direction using a Bayesian Decoder

The virtual head direction during sleep was estimated using a Bayesian decoding method (Peyrache et al., 2015; Zhang et al., 1998) that utilized the tuning curves of head direction cells recorded during wakefulness. This approach involves constructing probabilistic maps for each time bin (10ms, smoothed in 200-ms s.d. Gaussian windows for wake and REM, 50-ms windows for SWS), which represent the likelihood of each potential head direction based on the neural activity observed. The head direction for each time bin is then inferred by selecting the direction that maximizes this likelihood, assuming the neuronal firing follows a Poisson distribution. This Bayesian framework allows us to assess changes in internal HD during REM and SWS sleep. To estimate chance level decoding, we estimated decoding from time-inverted MEC spike trains.

During sleep, we estimated the decoded difference between the AD and MEC decoded directions. As a control, we subtracted MEC decoded direction based on time-inverted spikes from the ADn decoded direction.

### Evaluating ADn-MEC pairwise coordination with a GLM

To evaluate the degree of coordination between ADn and MEC, we implemented a Generalized Linear Model (GLM), following previous work (Gardner et al., 2019). We selected this approach because of the high synchronization observed in MEC unit activity during SWS. The GLM is particularly effective at disentangling the specific pairwise coupling from the influence of ongoing population activity.

Spike trains were binned into 40 ms intervals for wakefulness and REM sleep, and 10 ms intervals for SWS. Multi-unit activity (MUA) was calculated by summing the spiking activity of all MEC cells, excluding the neuron being modeled to avoid auto-correlation. The binned spike trains were then smoothed with a Gaussian kernel of 160ms for wake and REM epochs, and 10ms for SWS.

A Poisson GLM was used to model the firing rate of a single MEC neuron *r*_*MEC*_ in the MEC as a function of an individual ADn cell’s activity (*r*_*ADn*_) and the MEC population rate (*r*_*POP*_):

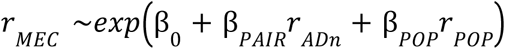

The coefficients β_*PAIR*_ quantifies the coupling between the ADn and MEC neurons, independently of fluctuation in population activity (β_*POP*_). The model intercept (β_0_) captures the baseline firing rate of the MEC neuron. GLMs were fitted using the *statsmodels* library (Seabold & Perktold, 2010).

To compute cross-correlograms of coupling strength, the predictors *r*_*ADn*_ and *r*_*POP*_ were time-shifted relative to the response variable, and the GLM was re-fit at each lag. This yielded a time-resolved coupling measure β_*PAIR*_ (t).

### Explaining MEC coupling based on ADn activity using a GLM

We sought to determine if the pairwise correlations in MEC could be explained by ADn activity, and vice versa. To investigate this, we extended the GLM approach described above to test for pairwise coupling within the same structure, once accounting for the input of a neuron in another structure:

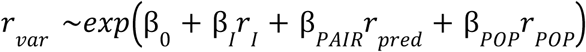

where *r*_*var*_, *r*_*I*_, *r*_*pred*_, *r*_*POP*_ are the binned firing rates of the target neuron (the *variable*), the input neuron (in another structure), the *predictor* neuron from the same structure, and the population from the same structure. The GLM yields the coefficients β that weight the contribution of each neuronal signal to the target neuron firing.

## ACKNOWLEDGMENTS

We would like to thank Lynda Mainville for technical support. We are thankful to members of the Peyrache laboratory for comments on the manuscript. This work was supported by the Canadian Research Chair in Systems Neuroscience (AP), CIHR Project Gransts 155957, 180330, and 190289 (AP), NSERC Discovery Grant RGPIN-2018-04600 (AP), Canada-Israel Health Research Initiative, jointly funded by the Canadian Institutes of Health Research, the Israel Science Foundation, the International Development Research Centre, Canada and the Azrieli Foundation 108877-001 (AP), the New Frontiers in Research Fund Exploration grant NFRFE-2021-00926 (AP).

## AUTHOR CONTRIBUTIONS

Conceptualization: GRV, AP; Methodology: GRV, AP; Software: GRV, AP; Validation: GV, AP; Formal analysis: GRV, AP; Investigation: GRV, MD, AP; Resources: AP; Data Curation: GRV; Writing: GRV, AP; Visualization: GRV, AP; Supervision: AP; Project administration: AP; Funding acquisition: AP.

## ADDITIONAL INFORMATION

Supplementary Information is available for this paper (Supplementary Figures 1-3).

## Supplementary Figures

**Figure S1.**
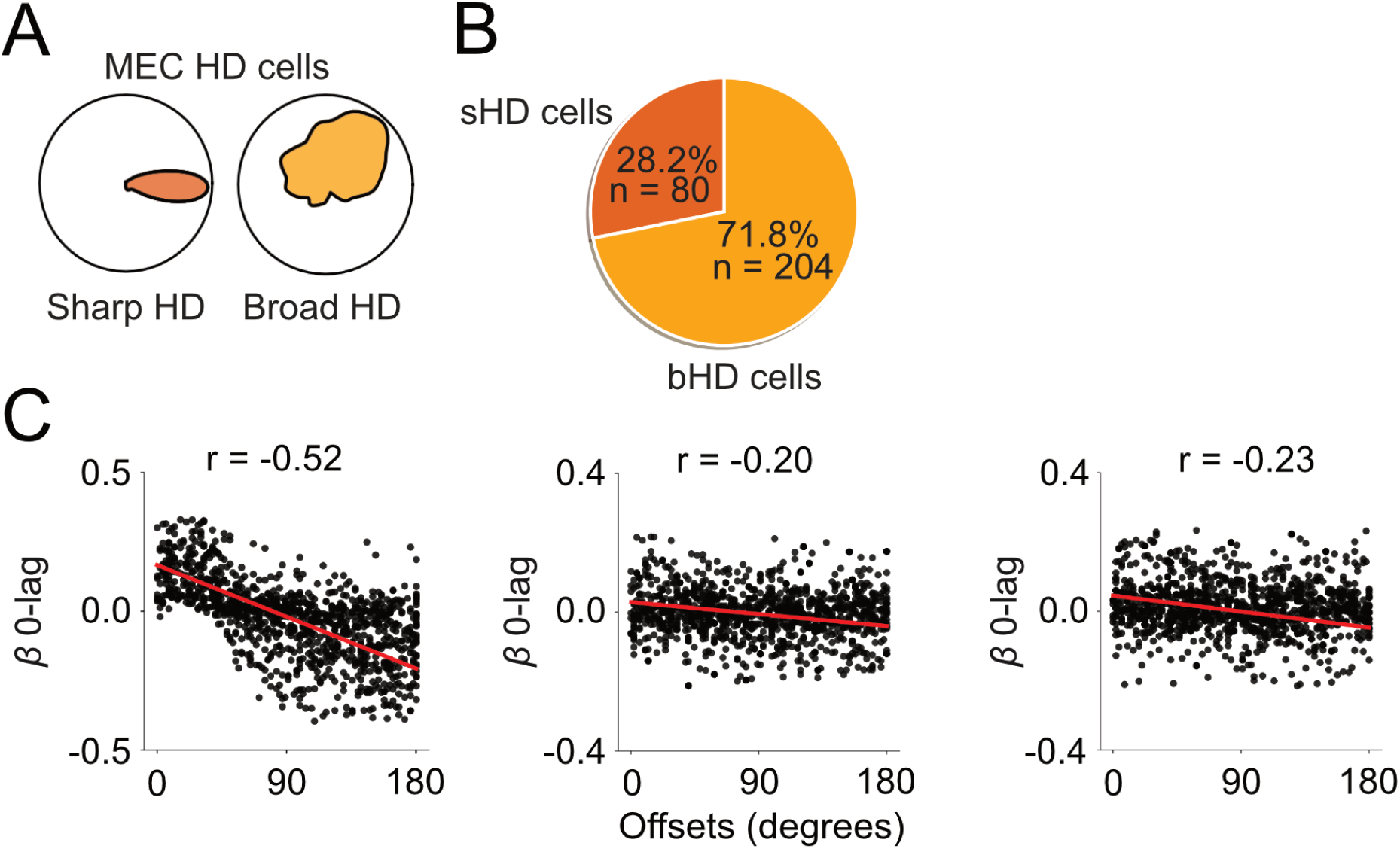
MEC broad HD cells have a low coupling with ADn during sleep. (A) Example of two HD cells classified as sharp and broad HD cells. Cells with a tuning curve larger than 90° were categorized as broad HD cells. (B) Proportion of cells classified as sharp HD cells (sHD, 28.2%) and broad HD cells (71.8%). (C) Beta values of 0-lag coupling plotted against the angular offset. *p* < 0.001 for all brain states.

**Figure S2.**
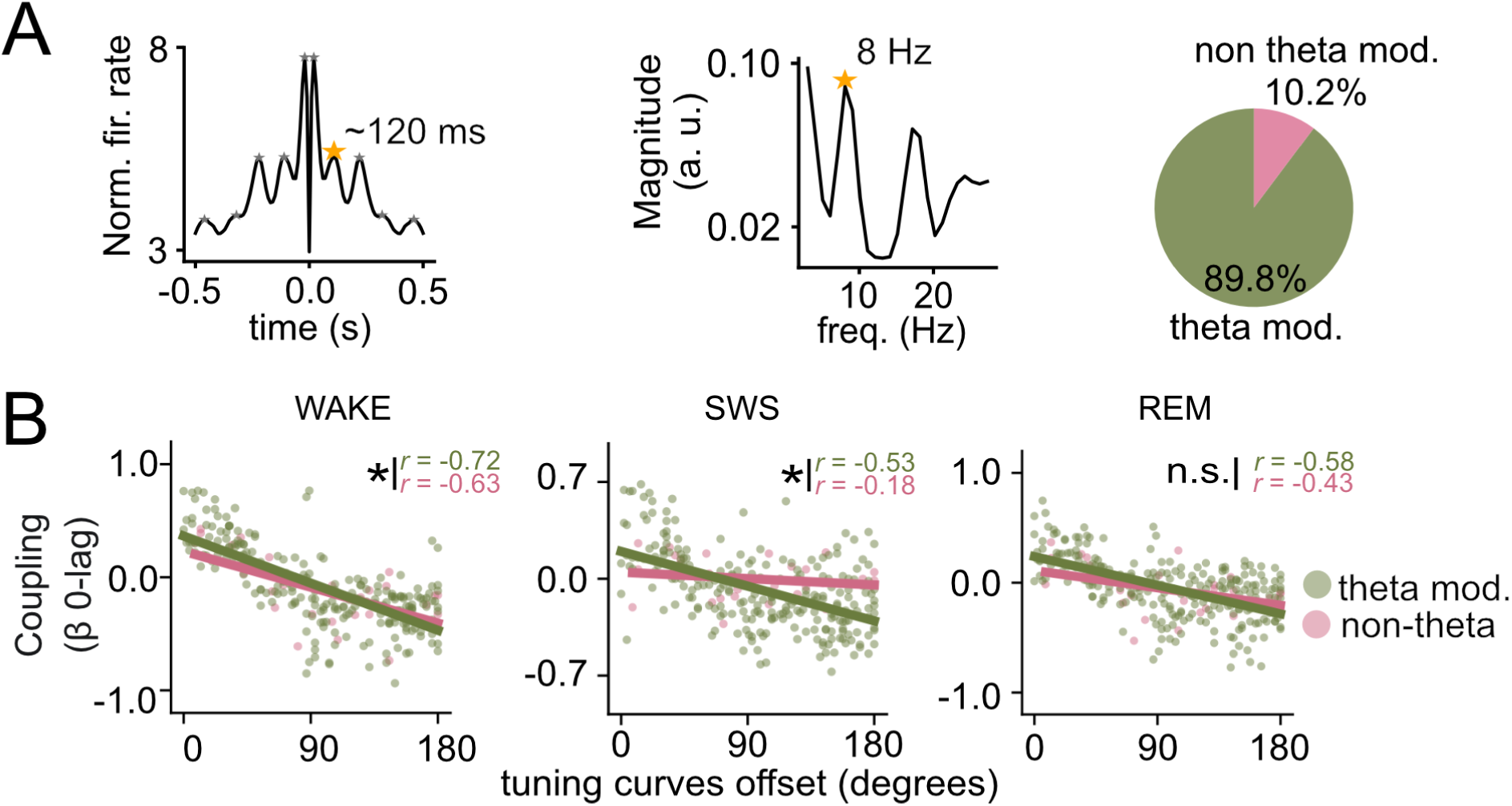
MEC non-theta-modulated HD cells lose their coupling with ADn during SWS. (A) From left to right, example of the autocorrelation of spikes from a theta rhythmic neuron recorded in MEC; fast Fourier transform from the autocorrelograms was used to detect oscillations at theta frequency (6-10 Hz); proportion of cells classified as theta-modulated (90%) and not theta-modulated (10%). (B) Beta values of 0-lag coupling plotted against the angular offset of the pairs color coded depending on theta-modulation. Insets, Pearson correlation *r* values (theta-modulated: p<0.001 (WAKE, SWS, REM); non-theta-modulated. Differences between correlations (theta vs. non-theta-modulated) were assessed using Fisher’s r-to-z transformation, followed by a z-test for independent correlation coefficients (α = 0.05).

